# A living vector field reveals constraints on galactose network induction in yeast

**DOI:** 10.1101/073817

**Authors:** Sarah R. Stockwell, Scott A. Rifkin

## Abstract

When a cell encounters a new environment, its transcriptional response can be constrained by its history. For example, yeast cells in galactose induce GAL genes with a speed and unanimity that depends on previous nutrient conditions. To investigate how cell-level gene expression dynamics produce population-level phenotypes, we built living vector fields from thousands of single-cell timecourses of the inducers Gal3p and Gal1p as cells switched to galactose from various nutrient histories. We show that, after sustained glucose exposure, the lack of GAL inducers leads to induction delays that are long but also variable; that cellular resources constrain induction; and that bimodally distributed expression levels arise from lineage selection -a subpopulation of cells induces more quickly and outcompetes the rest. Our results illuminate cellular memory in this important model system and illustrate how resources and randomness interact to shape the response of a population to a new environment.

**One Sentence Summary:** Single-cell galactose induction timecourses reveal that cellular resources and stochastic events determine which yeast cells outcompete their peers.

## Main Text

Budding yeast cells *(Saccharomyces cerevisiae)* can metabolize galactose by inducing a network of regulatory and metabolic genes, collectively known as the GAL genes (Fig. S1). When activated, the inducer Gal3p blocks the repressor Gal80p from inhibiting the action of the transcription factor Gal4p. Gal4p, in turn, promotes the transcription of the GAL genes, including the regulatory genes *GAL3, GAL1*, and *GAL80*, the membrane-bound galactose importer gene *GAL2*, and the enzymes *GAL1, GAL7*, and *GAL10* (1–5). The network's interlocking positive and negative regulatory feedback loops control induction in the presence of galactose (5–7). Abundant glucose represses GAL network activation (8–10).

The GAL network has been an important model system for metabolism, gene regulation, and now quantitative biology for most of a century, and the behavior of this network in various carbon sources at steady state is well understood (5–11). However, induction timecourses have revealed that the transient induction dynamics of the GAL network depend on cellular memory of previous nutrient environments (12). Cells previously grown in non-inducing/non-repressing media like raffinose or glycerol induce quickly and fairly uniformly (5, 13) (Fig. S2). The same is true for reinducing cultures: cells that have undergone prior galactose induction followed by short-term, 12-hour, glucose repression (14) before being switched back to pure galactose (Fig. 1; Fig. S2; Movie S1). By contrast, cell populations that have experienced long-term glucose repression (LTGR) induce the GAL genes after a long lag, producing a transiently bimodal distribution that, in population-level experiments, gradually resolves into an entirely induced population over the course of many hours (Fig. S2) (14, 15). The inducer/enzyme Gal1p is required for the reinduction phenotype (14), but the mechanisms and population biology behind the other memory phenotypes – particularly the transient bimodality after LTGR – are unknown, in part because research on GAL memory has largely been based on population-level measurements or snapshots of a population at a few times. As we show, such population-level measurements conflate the effects of growth and induction and mask the potential for competition between cell lineages to reshape the composition of the cell population (16).

**Fig. 1.**
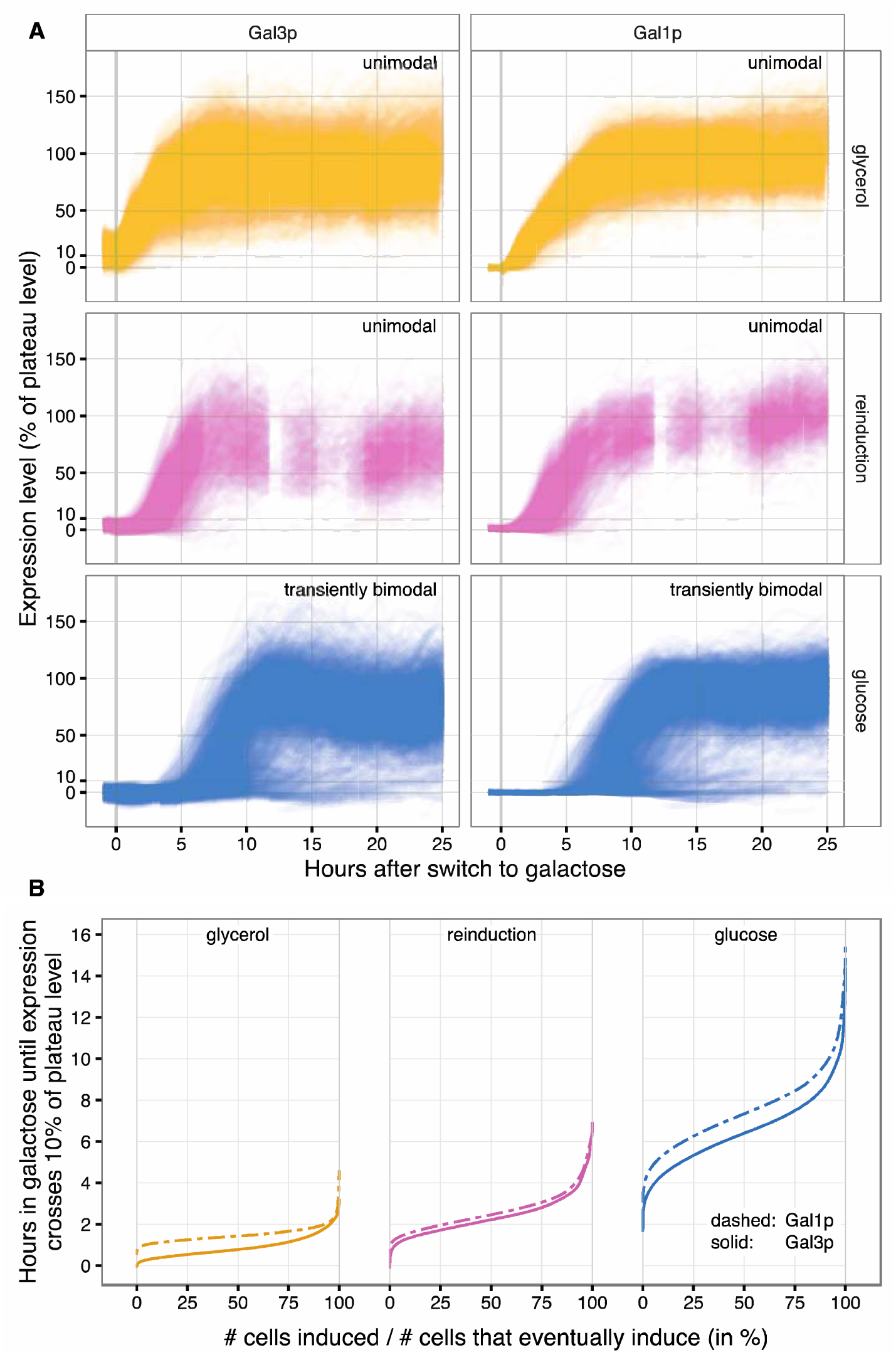
Cells induce quickly and uniformly after glycerol (yellow) and reinduction (pink), but variably after a long lag after long-term glucose repression (blue) producing a bimodal population distribution. (A) Single cell timecourses for each of the three history conditions for Gal3p-yECitrine and Gal1p-yECerulean. Expression levels have been normalized to an estimated 100% plateau level (see Methods). Each thin line corresponds to a single cell tracked over time, and each plot depicts trajectories from thousands of cells. Datapoints are spaced every 20 minutes and photobleaching is minimal (Fig S4). Absolute Gal1p-yECerulean fluorescence levels are approximately 10x higher than Gal3p-yECitrine, reflecting the massive induction of Gal1p. Fluorophore maturation time is expected to be on the order of 30 minutes. The y-axis represents percent of plateau-level gene expression (see Materials and Methods). 0% represents the median of the control (i.e. no fluorescent proteins) cell fluorescence levels for a given frame.(B) Empirical cumulative distributions for each history condition for Gal3p and Gal1p. Only cells that started with expression levels below 10% and induced to at least 75% of the plateau level are included (in the glucose condition in particular, most cells fail to induce).

To visualize the transient dynamics after the different media histories, we represent them as flows across a vector field on the state space of the two inducers, Gal1p and Gal3p (Fig. 2).Vector fields are standard modeling tools for analyzing dynamical systems. Here, we translated this tool into a biological reality, generating *living* vector fields to summarize our measurements of thousands of individual cells tracked over time (Fig. 1a) and giving us a comprehensive view of their induction dynamics. In these vector fields, each vector illustrates how Gal1p and Gal3p concentrations change over a given time interval. The root of a vector represents the protein concentrations at time Ti, the direction points towards the concentrations at time Ti+1, and the length is proportional to its speed. To measure Gal3p and Gal1p levels, we fused 2x-yECitrine to Gal3p and yECerulean to Gal1p (Fig. S3) and used a microfluidic device (17) to measure GAL network induction at 20-minute intervals (Fig. S4) as we switched cells with fluorescently labeled Gal1p and Gal3p proteins to 2% galactose from each of three conditions: glycerol-history, reinduction, and LTGR (Figs. S5-9; Materials and Methods).

**Fig. 2.**
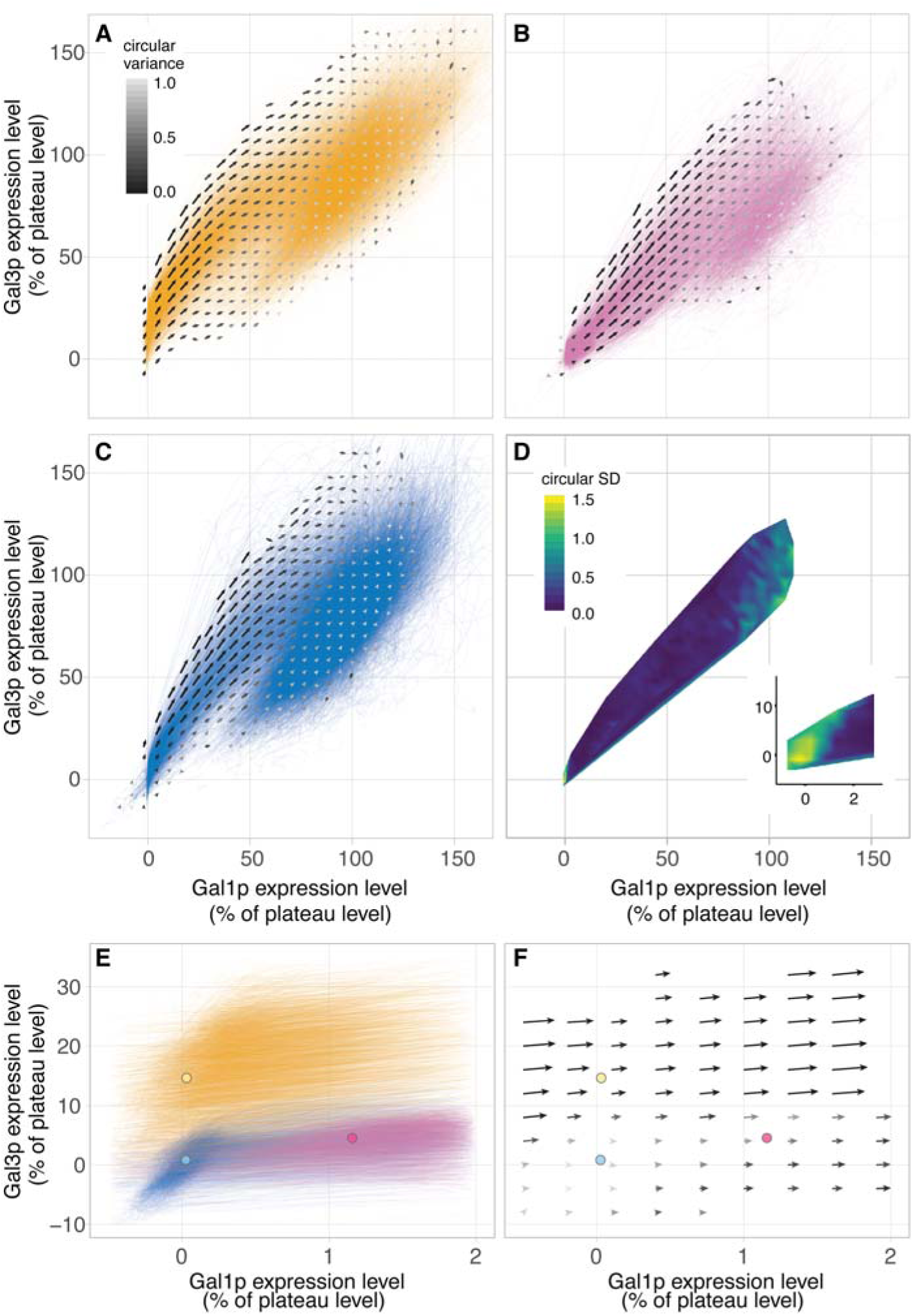
Empirical vector fields depicting the flow through the Gal3p/Gal1p state space for the three history conditions. (A-C) Vector fields for glycerol, reinduction, and long-term glucose repression conditions. Colored lines are traced of individual cells through the Gal3p/Gal1p state space. Arrows represent the vector field. The state space is binned in steps of 6% in each direction, and an arrow summarizes the movement of cells in its bin from one timepoint to the next. The length of the arrow is half the mean veolcity of cells, and the shade represents the circular variance (32) with black indicating consistency in the direction of displacement and white indicating inconsistency. The vector fields flow towards (100%,100%). (D) Consistency of the vector directions during induction in the three experiments. The color indicates the esimated circular standard deviation of the mean vector directions for each of the three experiments (see Methods). The inset shows the region at the corner near (0%,0%). Once cells leave this corner, the statistic drops to near zero indicating that cells are moving in a consistent direction regardless of experimental condition – they have lost their memories. (E) Cell trajectories near the (0%,0%) corner. Circles indicate the mean fluorescence values for each experiment at the time galactose was added. (F) The vector field near the (0%,0%) corner. Circles are as in (E).

The inducers Gal1p and Gal3p are not the only regulators of the GAL network, but they are among the most important. Gal3p has long been recognized as critical to induction speed (*gal3* mutants take days to induce the network instead of hours) (18–21), and Gal1p is responsible for fast, unimodal induction upon reinduction (14). The two inducers are linked by positive feedback loops (Fig. S1). In such a system, a cell beginning with low concentrations of the proteins would increase those proteins slowly, because neither inducer is abundant enough to ramp up GAL induction quickly. A cell with a moderate concentration of either protein will increase GAL expression quickly, because the positive feedback enables either inducer to accelerate the induction process for both. Finally, a cell approaching the equilibrium point where protein synthesis balances decay will begin to plateau and change protein concentrations slowly.

The vector field unifies the three memory phenotypes into one consistent picture of cell behavior. At steady state, glycerol cultures express detectable Gal3p but no Gal1p, and glucose cultures express neither (5). After 12 hours of glucose repression, reinduction cultures contain evident Gal1p (14) but little Gal3p (this study). The vector field representation illustrates how these three different initial inducer levels determine induction lag and population variability. Thus, each of the three history media places the cells different initial points in this state space (Figs. 2e–f). The appreciable presence of either inducer -Gal3p or Gal1p -is sufficient to drive the expression of the entire positive feedback loop, enabling these cells to ramp up induction quickly (Figs. 1a,b). However, cells that have been subjected to LTGR enter the galactose environment with no inducers to get the feedback loop going. They must wait for molecules of Gal3p to be produced so the feedback loop can begin to bootstrap the induction process. We suggest that this bootstrapping is the source of the long lag that has been observed following LTGR.

Single cell dynamics on the vector field could also explain the differences in population-level induction patterns (i.e. unimodal vs. transiently bimodal) among the three history conditions (14, 15) (Fig. 1a; Fig. S2). When only a few molecules of a protein are present in a cell, stochastic effects can dominate the molecular interactions involving that protein (22). LTGR-history cells are the only cases among our conditions where initial inducer concentrations are low enough that we might expect stochastic induction behavior. We hypothesize that when cells are switched to galactose after LTGR, they must wait for rare, stochastic molecular interactions to activate the positive feedback loops. As a result, individual cells would wait widely varying times before starting induction. This would produce a slow, sticky region of the Gal1p/Gal3p state space where the concentrations of both proteins would be near zero and from which cells would slowly escape one by one while they bootstrap themselves into GAL network expression. When extrapolated to the population level, this dynamic would manifest as the slow, transiently bimodal induction pattern characteristic of LTGR where an initial distribution of uninduced cells shifts to one that is completely induced. In contrast, in reinduction and glycerol-history conditions, cells start with appreciable levels of at least one inducer, placing them outside the putative sticky region. The dynamics in this fast deterministic regime would result in a unimodal induction pattern at the population level.

Population measurements (14, 15) cannot determine *why* the uninduced peak shrinks during the transiently bimodal period following LTGR. Does the uninduced fraction decrease because most of the cells in it activate the GAL network, thus switching to the induced fraction? Or does the fraction of uninduced cells shrink because a subpopulation of cells in it induces and starts to divide and demographically replace the rest? Our single-cell timecourses clearly illustrate the latter process: the fully-induced population is composed principally of the descendants of the earliest-inducing cells (Figs. 1a, 3a; Movies S1-2).

**Fig. 3.**
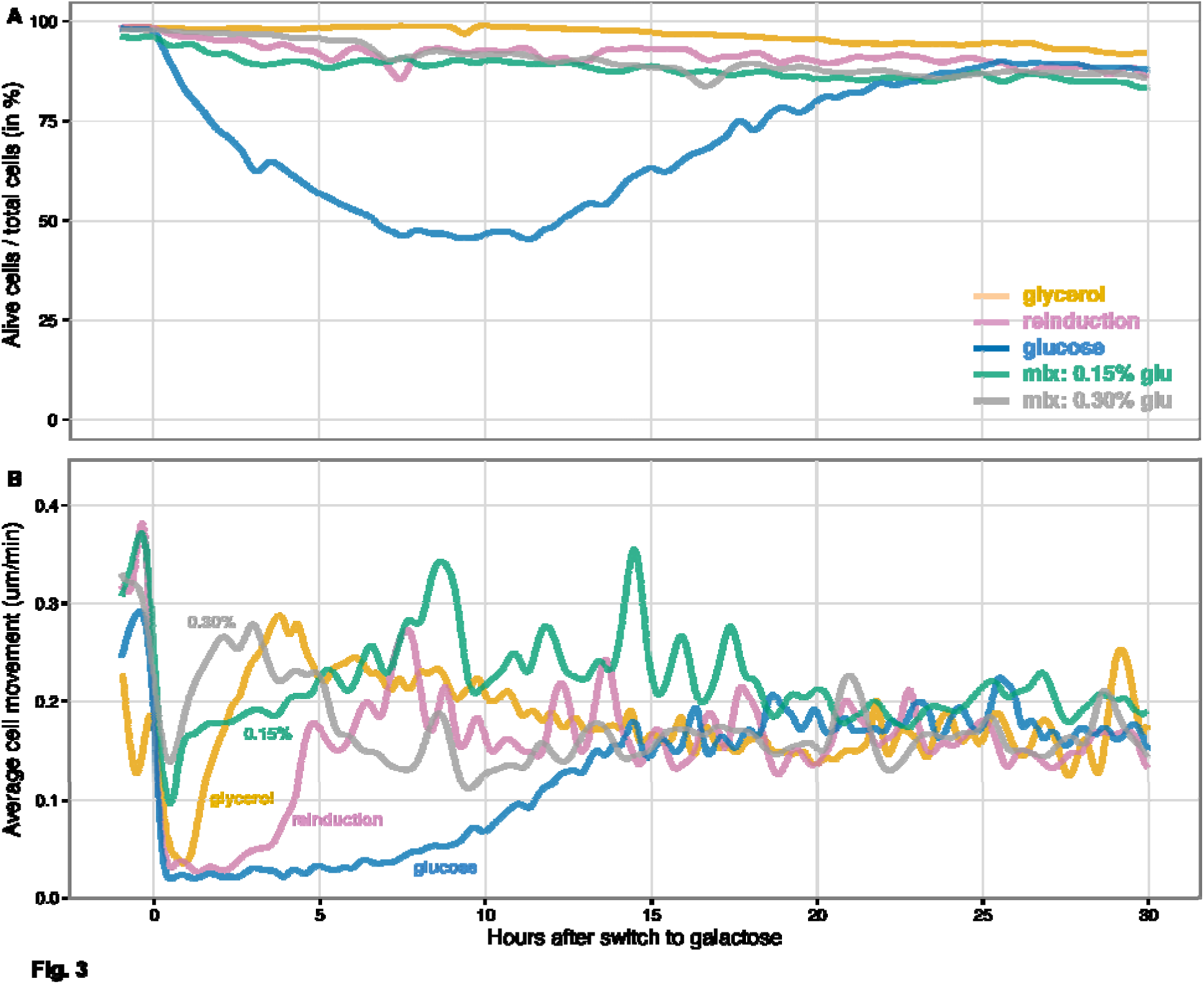
The switch to galactose imposes a heavy cost after long-term glucose repression. (A) An estimate of cell viability after the switch to galactose. Cells were classified as alive or dead (Materials and Methods; Fig. S8). Dying cells were classified as alive, so these curves represent upper bounds for the fraction of viable cells in the populations. (B) Average cell movement in microns per minute for cells in a field of view as estimated measuring physical displacements of individually tracked cells in bright field images taken every 2 minutes. Average cell movement is a surrogate for the amount of cell division since cells in the microfluidic device are confluent and push each other when they divide. After long-term glucose repression, the switch to galactose is accompanied by a 6 hour long pause in cell movement as cells bootstrap themselves into GAL network induction followed by a subsequent slow 7 hour recovery as the first cells to induce and their progeny take over the population (Movie S2).

The bootstrapping hypothesis makes a number of qualitative predictions for induction behavior. It suggests that cells starting in the region near (0%,0%) will have long and variable lag times and that the variability in lag times explains the bimodality of LTGR-history induction: as individual cells escape from this sticky region, they leave the uninduced population to join the inducing subpopulation. It also predicts that once cells leave the sticky region and accumulate appreciable levels of inducer, their induction trajectories will match those of cells in reinduction and glycerol-history conditions. We tested these predictions using the empirical vector fields derived from our microfluidics experiments.

The results of our microfluidics experiments fulfilled the predictions of the bootstrapping hypothesis (Fig. 1a). Lag times after LTGR were much longer and more variable than after the other two history conditions (Fig. 1b; Table S1), and many cells failed to induce. As a result of the long and variable lag, a small minority of cells induced, grew, and dominated the population, while others remained quiescent or died, producing the transiently bimodal induction distribution that others (14) have seen but not explained.

The small region around (0%, 0%) was populated almost exclusively by LTGR cells, and their initial trajectories differed markedly from those in other conditions (Fig. 2e–f). Once induction was underway, the dynamics were similar in all three histories (Fig. 2d; Fig. S11), which allowed us to map where the cells lose their memory of previous nutrient conditions (Fig. 2d). Cells moved in a consistent direction in the Gal3p/Gal1p state space and transitioned rapidly from low Gal3p to plateau levels of Gal3p (FigS. 1a,2. Their trajectories slowed as they approached plateau levels of Gal3p, and the flow on the vector field curved towards the fixed point (100%, 100%). The trajectories of Fig 1a show variation in expression levels for both proteins after most of the population has induced. In all three conditions, most of this variation can be explained by intercell differences in average expression level, as opposed to fluctuation in a cell’s measured expression over time (glycerol: 80.3%, 81.4% (Gal3p, Gal1p); reinduction: 86.2%, 87.4%; glucose: 80.1%, 84.5%; Materials and Methods). This suggests that while the trajectories of inducing cells are generally similar, their final expression plateaus are cell-specific and influenced by other variables.

Although the results from the microfluidics experiments support the bootstrapping hypothesis, it is possible that other factors could also contribute to lag time between history conditions. For example, glucose represses the expression of a large proportion of the yeast genome including the GAL genes and is known to suppress GAL expression via several different mechanisms (8, 10, 23–26). If glucose repression or its aftereffects linger in LTGR-history cells, this could cause the long lag we observe under those conditions.

Alternatively, resource constraints could contribute to induction delays. Gal1p is one of the most highly induced genes in yeast – it is upregulated 1000-fold – which makes the GAL1 promoter a useful tool for genetic engineering but also means that GAL network induction demands a large investment of resources (27, 28). Polymerases, ribosomes and carbon building blocks may need to be diverted from other genes, and a cell must have sufficient energy to build these proteins. Although cell populations with more energy reserves adapt as a whole more quickly to galactose (29, 30), cells do not use this battery power to propagate. Instead, when cells are switched from long-term glucose to galactose, every cell in the population abruptly stops growing or dividing and remains in stasis for hours (Fig. 3b; Movie S2). Although they are drowning in galactose, they have no GAL proteins and are unable to use it. Fueling initial GAL induction via stored energy/carbon reserves alone may be a formidable effort that only a few cells are able to muster (Fig. 3a). Only the cells that eventually begin GAL induction ever resume growth in our observations (Movie S1), and they rapidly outcompete the rest of the still-starving population, dooming them to demographic oblivion. The stately population-level picture of a bimodal population resolving into a unimodal, fully-induced one is, in fact, a process of lineage selection: some cells never manage to induce before they die, and others only do so too late. By contrast, the cultures that start with enough Gal3p or Gal1p to begin GAL induction quickly, and thus use galactose to fuel further GAL induction, have only a brief pause before growth resumes in the new medium (Fig. 3b), and the fully induced population preserves many of the original cell lineages (Fig. 1a; Movies S1-2).

If our resource-constraint hypothesis is correct, then adding a small amount of glucose to the galactose media should give cells a boost – extra resources to build their first GAL proteins (29). On the other hand, if lingering glucose repression were a cause of the induction delay, then glucose in the induction medium should prolong it. We found that adding glucose speeds up GAL network induction (Fig 4, Fig. S10): in both mixed-sugar conditions, half the inducing cells reach 10% of their plateau Gal3p expression levels within 2.1 hours (Fig. 4b), while LTGR cells take 6.4 hours to reach the same point (Fig. 1b). These results rule out lingering glucose repression as an explanation for the long lag times we observe after LTGR for times beyond 2.1 hours. They also point to energy or other resources as important limiting factors for the successful transition to metabolizing galactose. While the bootstrapping process imposes a lag on GAL activation when cells have no initial inducers, this process speeds up if cells can draw upon energy and carbon from the induction media (Fig. 4b). Without usable external energy, cells must fuel induction using only stored energy reserves and balance this against the need to live off these reserves in the meantime. Once GAL network induction is underway, galactose-derived resources become available to help fuel further induction (Fig. S12). Having other sources of energy available also gives cells the option whether the induce the GAL network at all (31).

**Fig. 4.**
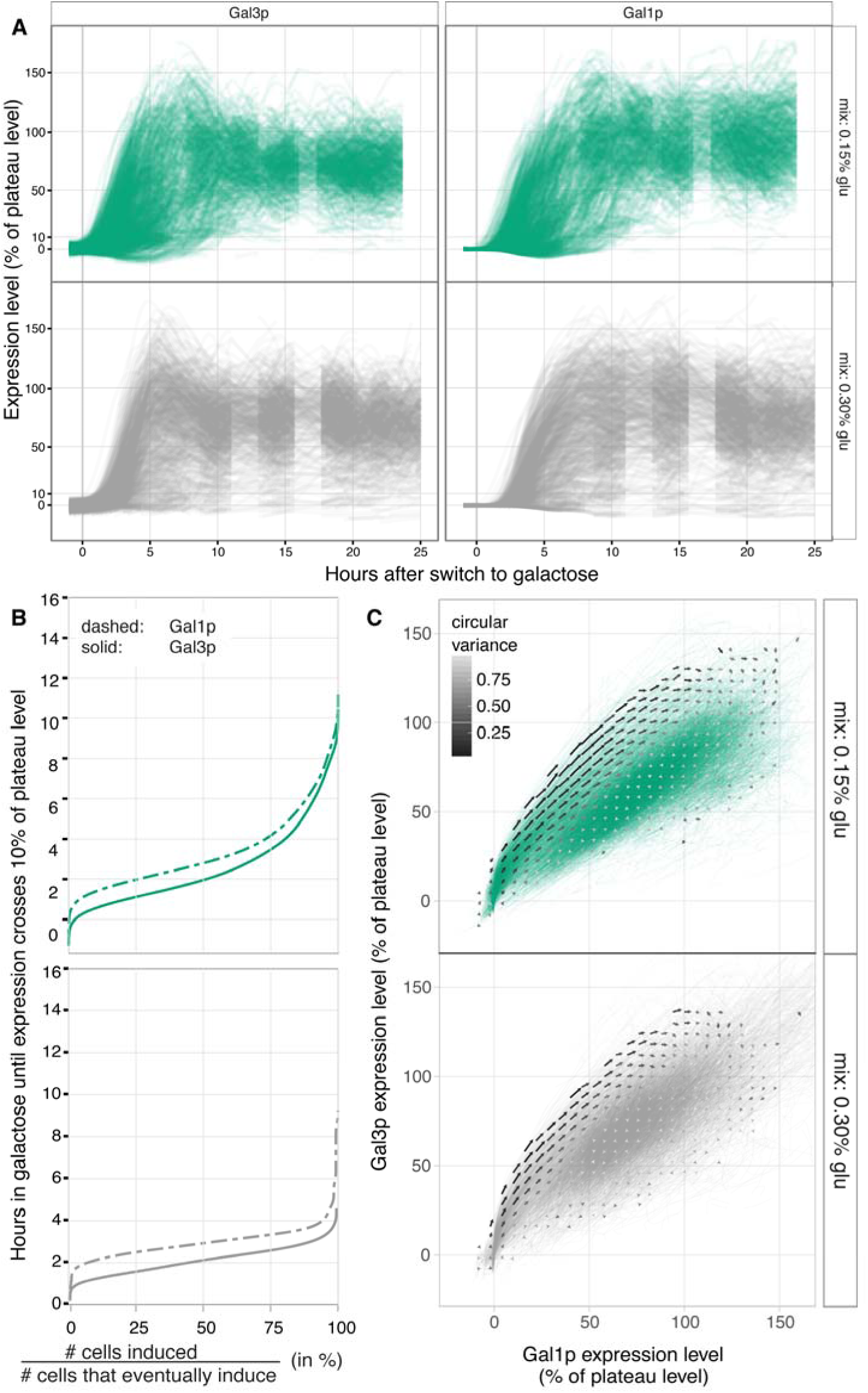
Adding a small amount of glucose speeds GAL network induction. (A) Single cell timecourses when cells are moved from long-term glucose repression to a mix of 0.15% glucose / 2% galatcose (green) or 0.3% glucose / 2% galatcose (grey). For both Gal3p and Gal1p, cells initially overshoot their long-term plateau levels. (B) Empirical cumulative distributions as in Fig. 1b. Additional glucose shortens the time until induction, but there is still appreciable between-cell variation with 0.15% glucose. (C) Vector fields as in Fig. 2a-c. The vector fields noticeably overshoot then curve back around towards (100%,100%). The flows on the Gal3p/Gal1p state space are similar in these mixes and otherwise similar to the conditions in Fig. 2.

Resource constraints alone, however, cannot explain the length and variabilty of the delay following LTGR. If they could, then the timing and variabilty of GAL network induction after long-term and short-term (reinduction) glucose repression should be the same. Or, if short-term exposure to rich glucose medium is not enough time to build up reserves, reinducing cells should take even longer to induce and lag length should be even more variable. In fact, induction after LTGR takes nearly three times longer than reinduction and is twice as variable (Fig. 1b; Table S1), so bootstrapping from initially absent inducers must play a role in lag times and variability. Conversely, if resource constraints played no role, then giving the cells extra resources during that critical period would have no effect. Instead, induction was faster when we supplemented cells with glucose. Therefore both bootstrapping and resource constraints affect the lag.

Although the *process* of bootstrapping after LTGR may be stochastic – because of the small numbers of molecules involved – it is entirely possible that *which* cells are successful, or perhaps the pool of cells that are able to induce at all, is influenced by aspects of cell state that we did not measure. When induction is easy and fast because of initially present inducers, there is no starvation period, and almost any cell can mount the effort required for induction. When induction is slow and requires surviving starvation, fewer cells have the energy or carbon reserves to last until they can bootstrap themselves to the induction level where they can eat galactose. Resource levels may also speed the bootstrapping process: when we supplemented cells with glucose after LTGR (Fig. 4), they induced more quickly.

When suddenly switched to galactose, yeast cells pay the cost of not being prepared. Activating the GAL network from a repressed state with no initial inducers requires resources that most cells do not muster and depends on chance molecular interactions that most cells do not experience. The minority of cell lineages that overcome these barriers take over the population, The living vector fields synthesize results from five different nutrient conditions to reveal that GAL induction behavior depends on whether cells need to bootstrap the positive feedback loops in the network from initially absent inducers and that the bootstrapping process is constrained by cellular resources.

## Acknowledgments

This work was supported by an IRACDA postdoctoral fellowship to SRS (NIH K12 GM068524), a Human Frontiers Science Program Young Investigator award to SAR (HFSP RGY0073/2010), and the San Diego Center for Systems Biology (NIH P50 GM085764). We thank Christian Landry, Mike Ferry, Ivan Razinkov, Jeff Hasty, and members of the UCSD Microscopy and Modeling group for technical help, discussion, and suggestions.

## References and Notes

1. E. Giniger, M. Ptashne, Cooperative DNA binding of the yeast transcriptional activator GAL4. Proc Natl Acad Sci U S A. 85, 382–6 (1988).

2. G. O. Bryant, M. Ptashne, Independent recruitment in vivo by Gal4 of two complexes required for transcription. Mol Cell. 11, 1301–9 (2003).

3. R. J. Bram, N. F. Lue, R. D. Kornberg, A GAL family of upstream activating sequences in yeast: roles in both induction and repression of transcription. EMBO J. 5, 603–8 (1986).

4. D. Abramczyk, S. Holden, C. J. Page, R. J. Reece, Interplay of a ligand sensor and an enzyme in controlling expression of the Saccharomyces cerevisiae GAL genes. Eukaryot Cell. 11, 334–42 (2012).

5. D. Lohr, P. Venkov, J. Zlatanova, Transcriptional regulation in the yeast GAL gene family: a complex genetic network. FASEB J. 9, 777–87 (1995).

6. S. A. Ramsey, J. J. Smith, D. Orrell, M. Marelli, T. W. Petersen, P. de Atauri, H. Bolouri, J. D. Aitchison, Dual feedback loops in the GAL regulon suppress cellular heterogeneity in yeast. Nat Genet. 38, 1082–7 (2006).

7. M. Acar, A. Becskei, A. van Oudenaarden, Enhancement of cellular memory by reducing stochastic transitions. Nature. 435, 228–32 (2005).

8. M. Johnston, M. Carlson, in The Molecular and Cellular Biology of the Yeast Saccharomyces: Gene Expression, E. Jones, J. Pringle, J. Broach, Eds. (Cold Spring Harbor Laboratory Press, Cold Spring Harbor, NY, 1992), pp. 193–281.

9. G. O. Bryant, V. Prabhu, M. Floer, X. Wang, D. Spagna, D. Schreiber, M. Ptashne, Activator control of nucleosome occupancy in activation and repression of transcription. PLoS Biol. 6, 2928–39 (2008).

10. J. O. Nehlin, M. Carlberg, H. Ronne, Control of yeast GAL genes by MIG1 repressor: a transcriptional cascade in the glucose response. EMBO J. 10, 3373–7 (1991).

11. E. Braun, N. Brenner, Transient responses and adaptation to steady state in a eukaryotic gene regulation system. Phys Biol. 1, 67–76 (2004).

12. S. R. Stockwell, C. R. Landry, S. A. Rifkin, The yeast galactose network as a quantitative model for cellular memory. Mol Biosyst. 11, 28–37 (2015).

13. M. Johnston, J. S. Flick, T. Pexton, Multiple mechanisms provide rapid and stringent glucose repression of GAL gene expression in Saccharomyces cerevisiae. Mol Cell Biol. 14, 3834–41 (1994).

14. I. Zacharioudakis, T. Gligoris, D. Tzamarias, A yeast catabolic enzyme controls transcriptional memory. Curr Biol. 17, 2041–6 (2007).

15. S. R. Biggar, G. R. Crabtree, Cell signaling can direct either binary or graded transcriptional responses. EMBO J. 20, 3167–76 (2001).

16. D. Nevozhay, R. M. Adams, E. Van Itallie, M. R. Bennett, G. Balázsi, Mapping the Environmental Fitness Landscape of a Synthetic Gene Circuit. PLoS Comput Biol. 8, e1002480 (2012).

17. M. S. Ferry, I. A. Razinkov, J. Hasty, in Methods in Enzymology, Chris Voigt, Ed. (Academic Press, 2011; http://www.sciencedirect.com/science/article/pii/B9780123850751000147), vol. Volume 497, pp. 295–372.

18. H. C. Douglas, G. Pelroy, A gene controlling inducibility of the galactose pathway enzymes in Saccharomyces. Biochimica et biophysica acta. 68, 155–156 (1963).

19. S. Spiegelman, R. R. Sussman, E. Pinska, On the Cytoplasmic Nature of “Long-Term Adaptation” in Yeast. Proc Natl Acad Sci U S A. 36, 591–606 (1950).

20. P. J. Bhat, J. E. Hopper, The mechanism of inducer formation in gal3 mutants of the yeast galactose system is independent of normal galactose metabolism and mitochondrial respiratory function. Genetics. 128, 233–9 (1991).

21. O. Winge, C. Roberts, Inheritance of enzymatic characters in yeasts, and the phenomenon of long-term adaptation. Compt. Rend. de Labor. Carlsberg. Ser. Physiol. 24, 263–315 (1948).

22. H. H. McAdams, A. Arkin, Stochastic mechanisms in gene expression. PNAS. 94, 814–819 (1997).

23. M. Carlson, Glucose repression in yeast. Curr Opin Microbiol. 2, 202–7 (1999).

24. B. G. Adams, Induction of galactokinase in Saccharomyces cerevisiae: kinetics of induction and glucose effects. J Bacteriol. 111, 308–15 (1972).

25. D. W. Griggs, M. Johnston, Regulated expression of the GAL4 activator gene in yeast provides a sensitive genetic switch for glucose repression. Proc Natl Acad Sci U S A. 88, 8597–601 (1991).

26. M. S. Lamphier, M. Ptashne, Multiple mechanisms mediate glucose repression of the yeast GAL1 gene. Proc Natl Acad Sci USA. 89, 5922–6 (1992).

27. M. Johnston, A model fungal gene regulatory mechanism: the GAL genes of Saccharomyces cerevisiae. Microbiol Rev. 51, 458–76 (1987).

28. B. L. Baumgartner, M. R. Bennett, M. Ferry, T. L. Johnson, L. S. Tsimring, J. Hasty, Antagonistic gene transcripts regulate adaptation to new growth environments. Proc Natl Acad Sci USA. 108, 21087–92 (2011).

29. S. Spiegelman, J. M. Reiner, R. Cohnberg, The relation of enzymatic adaptation to the metabolism of endogenous and exogenous substrates. J Gen Physiol. 31, 27–49 (1947).

30. J. M. Reiner, S. Spiegelman, The preadaptive oxidation of galactose by yeast. J Gen Physiol. 31, 51–74 (1947).

31. R. Escalante-Chong, Y. Savir, S. M. Carroll, J. B. Ingraham, J. Wang, C. J. Marx, M. Springer, Galactose metabolic genes in yeast respond to a ratio of galactose and glucose. PNAS. 112, 1636–1641 (2015).

32. A. Pewsey, M. Neuhäuser, G. D. Ruxton, Circular statistics in R (Oxford University Press, Oxford, 2013).

33. C. Baker Brachmann, A. Davies, G. J. Cost, E. Caputo, J. Li, P. Hieter, J. D. Boeke, Designer deletion strains derived from Saccharomyces cerevisiae S288C: A useful set of strains and plasmids for PCR-mediated gene disruption and other applications. Yeast. 14, 115–132 (1998).

34. M. A. Sheff, K. S. Thorn, Optimized cassettes for fluorescent protein tagging in Saccharomyces cerevisiae. Yeast. 21, 661–670 (2004).

35. N. C. Shaner, R. E. Campbell, P. A. Steinbach, B. N. G. Giepmans, A. E. Palmer, R. Y. Tsien, Improved monomeric red, orange and yellow fluorescent proteins derived from Discosoma sp. red fluorescent protein. Nat Biotech. 22, 1567–1572 (2004).

36. A. D. Edelstein, M. A. Tsuchida, N. Amodaj, H. Pinkard, R. D. Vale, N. Stuurman, Advanced methods of microscope control using μManager software. Journal of Biological Methods. 1, 10 (2014).

37. The MathWorks, Matlab (Natick, MA, 2009).

38. M. Ricicova, M. Hamidi, A. Quiring, A. Niemistö, E. Emberly, C. L. Hansen, Dissecting genealogy and cell cycle as sources of cell-to-cell variability in MAPK signaling using high-throughput lineage tracking. PNAS. 110, 11403–11408 (2013).

39. N. Otsu, A Threshold Selection Method from Gray-Level Histograms. IEEE Transactions on Systems, Man, and Cybernetics. 9, 62–66 (1979).

40. H. W. Kuhn, The Hungarian method for the assignment problem. Naval research logistics quarterly. 2, 83–97 (1955).

41. L. Breiman, Random forests. Mach Learn. 45, 5–32 (2001).

42. K. V. Mardia, Statistics of directional data: probability and mathematical statistics (Academic Press, London, 1972).

43. MySQL (www.mysql.org).

44. R Core Team, R: A Language and Environment for Statistical Computing (R Foundation for Statistical Computing, Vienna, Austria, 2016; https://www.R-project.org/).

45. H. Wickham, R. Francois, dplyr: A Grammar of Data Manipulation (2015; https://CRAN.R-project.org/package=dplyr).

46. H. Wickham, The Split-Apply-Combine Strategy for Data Analysis. Journal of Statistical Software. 40, 1–29 (2011).

47. RStudio Team, RStudio: Integrated Development Environment for R (RStudio, Inc., Boston, MA, 2015; http://www.rstudio.com/).

48. M. Tsagris, G. Athineou, Directional: Directional Statistics (2016; https://CRAN.R-project.org/package=Directional).

49. D. Bates, M. Mächler, B. Bolker, S. Walker, Fitting Linear Mixed-Effects Models Using lme4. Journal of Statistical Software. 67, 1–48 (2015).

50. H. Wickham, ggplot2: Elegant Graphics for Data Analysis (Springer-Verlag New York, 2009; http://ggplot2.org).

